# The microbial basis of impaired wound healing: differential roles for pathogens, “bystanders”, and strain-level diversification in clinical outcomes

**DOI:** 10.1101/427567

**Authors:** Lindsay Kalan, Jacquelyn S. Meisel, Michael A. Loesche, Joseph Horwinski, Ioana Soaita, Xiaoxuan Chen, Sue E. Gardner, Elizabeth A. Grice

## Abstract

Chronic, non-healing wounds are a major complication of diabetes associated with high morbidity and health care expenditures estimated at $9-13 billion annually in the US. Though microbial infection and critical colonization is hypothesized to impair healing and contribute to severe outcomes such as amputation, antimicrobial therapy is inefficacious and the role of microbes in tissue repair, regeneration, and healing remains unclear. Here, in a longitudinal prospective cohort study of 100 subjects with non-infected neuropathic diabetic foot ulcer (DFU), we performed metagenomic shotgun sequencing to elucidate microbial temporal dynamics at strain-level resolution, to investigate pathogenicity and virulence of the DFU microbiome with respect to outcomes, and to determine the influence of therapeutic intervention on the DFU microbiota. Slow healing DFUs were associated with signatures of biofilm formation, host invasion, and virulence. Though antibiotic resistance was widespread at the genetic level, debridement, rather than antibiotic treatment, significantly shifted the DFU microbiome in patients with more favorable outcomes. Primary clinical isolates of *S. aureus, C. striatum*, and *A. faecalis* induced differential biological responses in keratinocytes and in a murine model of diabetic wound healing, with the *S. aureus* strain associated with non-healing wounds eliciting the most severe phenotype. Together these findings implicate strain-level diversification of the wound pathogen *S. aureus* in chronic wound outcomes, while revealing potential contributions from skin commensals and other previously underappreciated constituents of the wound microbiota.

## INTRODUCTION

Chronic, non-healing wounds are common and costly complications of diabetes. Up to one in four persons with diabetes will develop a diabetic foot ulcer (DFU)(Boyko et al., 2015; Martins-Mendes et al., 2014), and approximately 25% of hospital stays for patients with diabetes are due to infected or ischemic DFU(Ramsey et al., 1999). Complications from DFUs account for two-thirds of all non-traumatic lower extremity amputations performed in the United States(Hoffstad et al., 2015; Martins-Mendes et al., 2014) and 5-year mortality rates surpass those of prostate and breast cancer, among others(Armstrong et al., 2007; Moulik et al., 2003). Improved therapeutic approaches are desperately needed, as morbidity, mortality, and health care expenditures only continue to increase as the prevalence of diabetes escalates worldwide.

Microbial colonization, biofilm formation, and infection are hypothesized to impair healing of DFUs and contribute to severe complications such as osteomyelitis and amputation. Wound infection is believed to underlie up to 90% of amputations(Boulton et al., 2005); yet quantitative cultures of uninfected DFUs were not predictive of outcomes(Gardner et al., 2014). Systemic and topical antimicrobials are often used to treat DFUs, despite their limited efficacy and even though it is often unclear which microorganisms are pathogenic and if some microorganisms may confer a beneficial effect. Culture-based methods, which are biased toward those microorganisms that thrive under laboratory conditions, insufficiently represent fungal and bacterial communities that colonize DFUs and other chronic wounds(Gardner et al., 2013). The role of microbial bioburden in DFU outcomes and complications remains unclear, including the significance of microbial load and diversity and the role of specific microorganisms including known wound pathogens and microorganisms considered as skin commensals or environmental contaminants.

Culture-independent, amplicon-based sequencing methods (i.e. bacterial and fungal ribosomal RNA gene sequencing) have highlighted the polymicrobial and temporally dynamic nature of the bacterial and fungal microbiota colonizing DFU. However, only limited insight has been gained with these methods regarding the role of wound microbiota in patient outcomes, complications, and healing(Kalan et al., 2016; Loesche et al., 2017). A major limitation of such approaches is the poor taxonomic resolution that precludes accurate identification to the species or strain level(Meisel et al., 2016). Mounting evidence suggests that genetically distinct strains within a single species have important functional differences that influence interactions with their host(Byrd et al., 2017). Shotgun metagenomics, the untargeted sequencing of bulk microbial genomes in a specimen, could address this limitation while providing insight into the functions and virulence of the DFU microbiota. While technically and computationally challenging when applied to clinical wound specimens that contain abundant “contaminating” human tissues and cells, shotgun metagenomics has the potential for unprecedented insight into the microbial basis of impaired wound healing while revealing clinically important biomarkers of healing and complication. These biomarkers can then be combined with other individual and contextual factors to identify and target subgroups of patients for prevention and treatment, consistent with the evolving view and potential of precision health(Whitson et al., 2016).

For these reasons, we performed shotgun metagenomic sequencing of DFU samples to identify strain-level diversity and to profile the genomic content of the DFU microbiota. We identified features of the DFU microbiome, including individual strains and pathogenicity/virulence factors, that are associated with poor healing and outcomes. Using cultured clinical isolates from the same DFU patients, we show that microbes previously labeled as skin commensals, wound pathogens, and environmental contaminants have differential and strain-specific effects on the host that influence cytokine production by human keratinocytes and wound closure in a murine model of diabetic wound healing.

## RESULTS

### Overview of study cohort and design

We enrolled 100 subjects with neuropathic, plantar DFU to examine the relationship between wound bioburden and clinical outcome. All enrolled subjects were free of clinical signs of infection at presentation and free from antibiotic exposure for >2 weeks. Specimens were obtained from DFUs by Levine’s swab (Fig. 1A), which samples the deep tissue fluid. Clinical factors were concurrently measured and recorded, including: blood glucose control (total blood glucose; hemoglobin A1c, HgbA1c), inflammation (white blood cell count, WBC; C reactive protein, CRP), ischemia (ankle-brachial index, ABI; toe-brachial index, TBPI), and wound oxygen levels (transcutaneous oxygen pressure at the wound edge). All patients underwent aggressive surgical debridement immediately following the first wound specimen collection at t=0. Specimens were obtained every two weeks, following conservative sharp debridement and non-bacteriostatic saline cleansing, until the wound healed, resulted in an amputation, or remained unhealed at the end of the 26-week follow-up period. After filtering out patients with wounds that healed before the collection of the first time point (t=2 weeks), dropped from the study for unknown reasons or due to another infection (e.g., respiratory infection), or had missing samples, we proceeded with shotgun metagenomic sequencing to result in 195 reconstructed metagenomes from 46 patient timelines. A detailed description of the clinical co-variates is provided in Table 1. Complications were experienced by 17 (37%) of the 46 subjects defined as: 1) wound deterioration, 2) development of osteomyelitis, and/or 3) amputations.

**Table 1.**
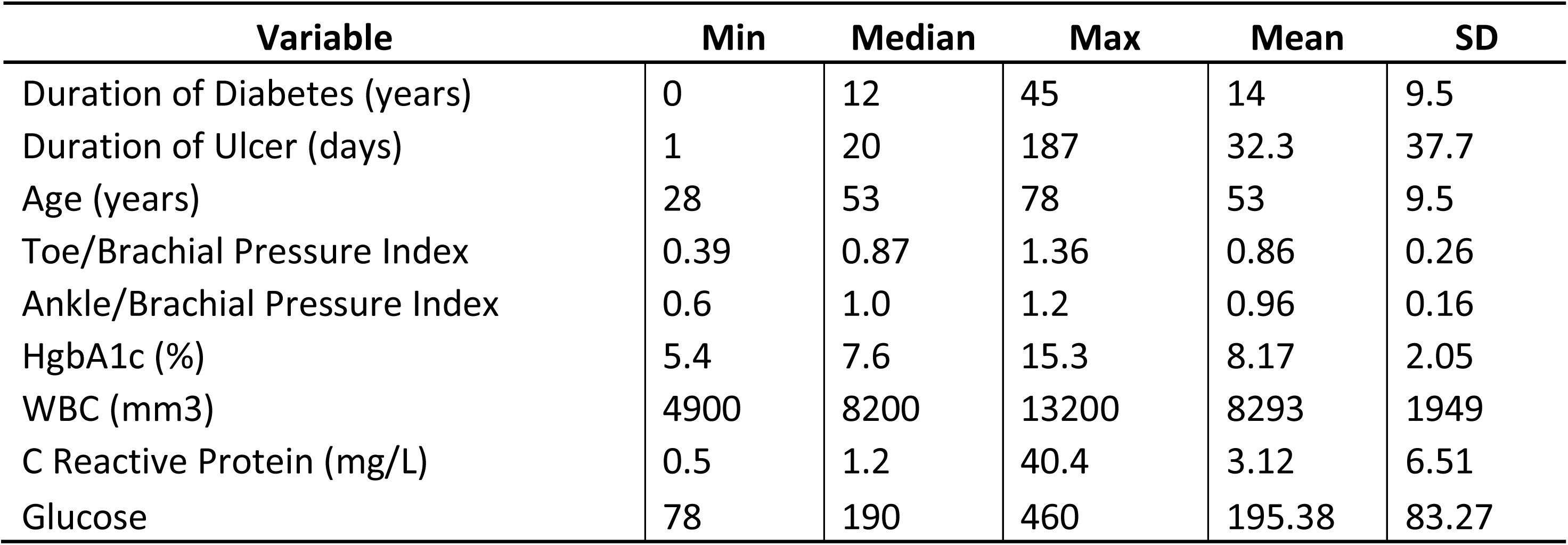
Summary Statistics for Clinical Variables at the Baseline Visit (n=46)

**Figure 1:**
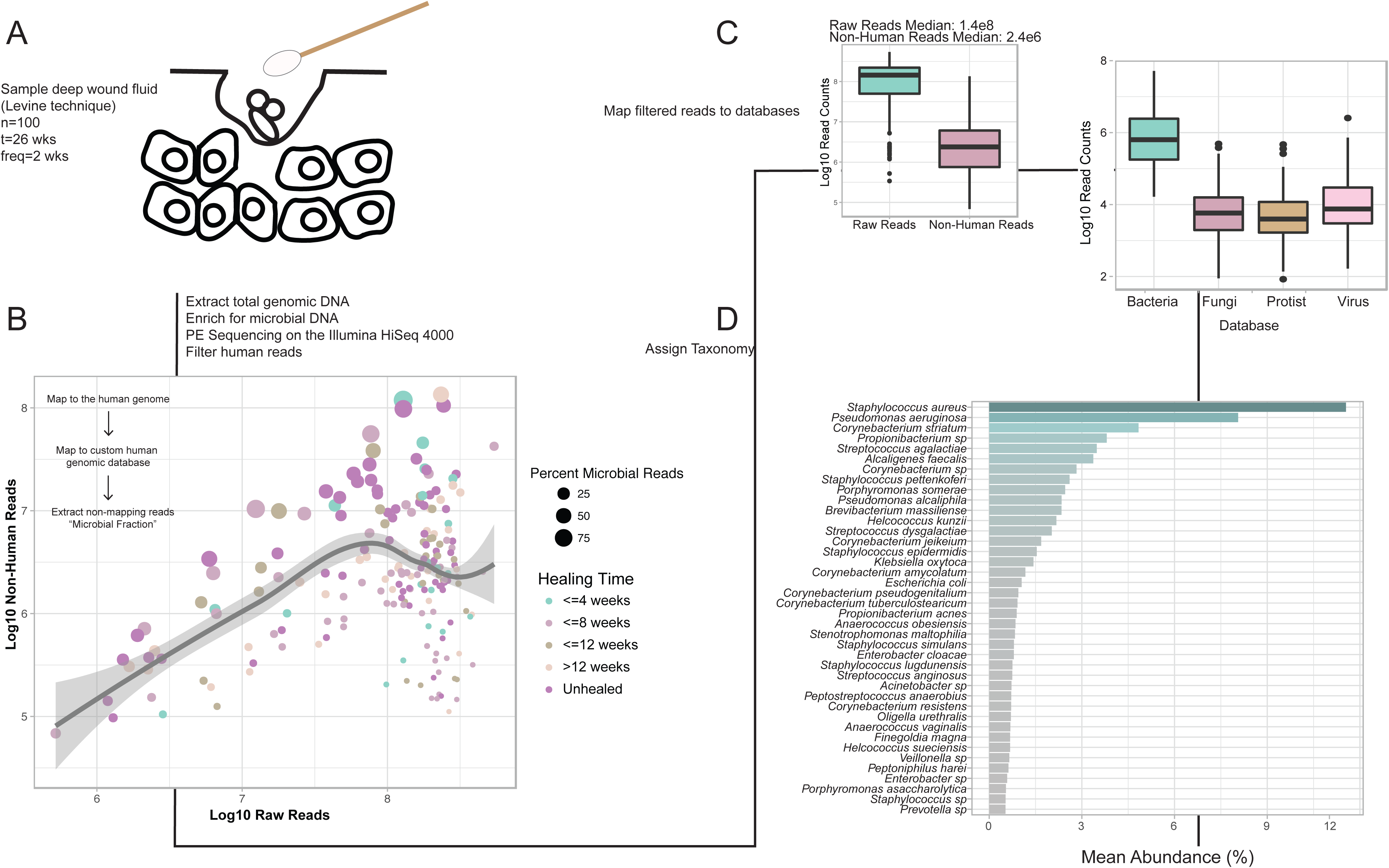
Shotgun metagenomic sequencing of the diabetic foot ulcer microbiome. A) The Levine technique(Levine et al., 1976) was used to sample deep wound fluid from ulcers every two weeks over a period of 26 weeks (n=46 subjects). Microbial DNA was enriched from samples by bead based eukaryotic DNA depletion prior to whole shotgun metagenome sequencing (n=195 samples). B) Reads mapping to the human genome and a custom database of human sequences were filtered prior to analysis. Increasing sequencing depth results in a linear increase in the fraction of total microbial reads. C) Non-human reads are mapped to phylogeny-based bacterial, fungal, protist, and viral databases for classification. D) Mean abundance of bacterial species detected in >0.5% abundance of all samples and at least 1% abundance in individual samples.

### Diversity and composition of the DFU metagenome and concordance with 16S rRNA gene amplicon data

We obtained a median of 144,416,914 reads per sample, and microbial reads comprised 0.04% to 92.55% of raw sequence reads (median = 2.52%). Increasing sequence depth increased the number of microbial reads linearly until saturation occurred at approximately 1×10^8^ reads (Fig. 1B). After filtering reads mapping to human genome references, the median number of microbial reads was 2,381,624 reads per sample (Fig. 1C). After mapping reads to multi-kingdom reference databases, bacterial reads comprised the largest proportion of microbial reads detected (96%), with *Staphylococcus aureus, Pseudomonas aeruginosa, Corynebacterium striatum*, and *Alcaligenes faecalis*, respectively, comprising the most abundant species of bacteria detected in all samples (Fig. 1C-D).

We assessed the concordance of shotgun metagenomic sequencing with 16S rRNA gene sequencing as previously reported for this same cohort(Loesche et al., 2017). Previous culture-independent studies of wound microbiota have all used amplicon-based sequencing approaches, which have poor concordance with culture-based measures of bioburden(Gardner et al., 2013, 2014; Rhoads et al., 2012). We assessed the concordance between shotgun metagenomic and 16S rRNA gene amplicon sequencing using two alpha diversity metrics (Supplementary Fig. 1). Shannon diversity, a measure of species richness and evenness within a sample, was concordant between our two datasets (ρ=0.36; P≤0.0001). Species richness, measured by the number of genus-level operational taxonomic units (OTU richness; 16S rRNA amplicon sequencing) or genera (shotgun metagenomics) detected per sample, was also concordant (ρ=0.22; P≤0.01) (Supplementary Fig. 1).

The most abundant genera in our data set were, in descending order, *Staphylococcus* (18.95%), *Corynebacterium* (14.64%), *Pseudomonas* (9.37%), and *Streptococcus* (7.32%) (Fig. 2A). These genera are consistent with 16S rRNA gene amplicon data but further highlight the limitation of taxonomic resolution of current marker gene approaches (Supplementary Fig. 2A). *Staphylococcus* and *Corynebacterium* are genera that encompass both species that are normal constituents of the healthy skin environment (e.g., *S. epidermidis* or *C. amycolatum*) and species associated with pathogenesis (e.g., *S. aureus or C. striatum*). Shotgun metagenomics revealed that *S. aureus* was the major *Staphylococcus* species identified, and was dominated by a single strain, *S. aureus* 7372. Staphylococcal species present in lesser abundance included the coagulase negative species *S. pettenkoferi*, *S. epidermidis, S. simulans*, and *S. lugdunensis. Corynebacterium striatum*, a bacterium associated with infection and multi-drug resistance(G.J. et al., 2010; Hahn et al., 2016; Patel et al., 2016), was the most prevalent *Corynebacterium* spp. classified in DFU, with *C. jeikeium, C. amycolatum, C. pseudogenitalium, C. tuberculostearicum*, and *C. resistens* in lesser abundances (Fig. 2B). *Pseudomonas* spp. were the third most abundant genera detected with the most abundant species identified as *P. aeruginosa* followed by *P. alcaliphila. P. aeruginosa* is regarded as a common pathogen associated with DFU as it is frequently isolated by culture-based methods (Supplementary Fig. 2B). *Streptococcus* was the fourth most abundant genera, with *S. agalactiae, S. dysgalactiae*, and *S. anginosus* present in descending abundance.

**Figure 2:**
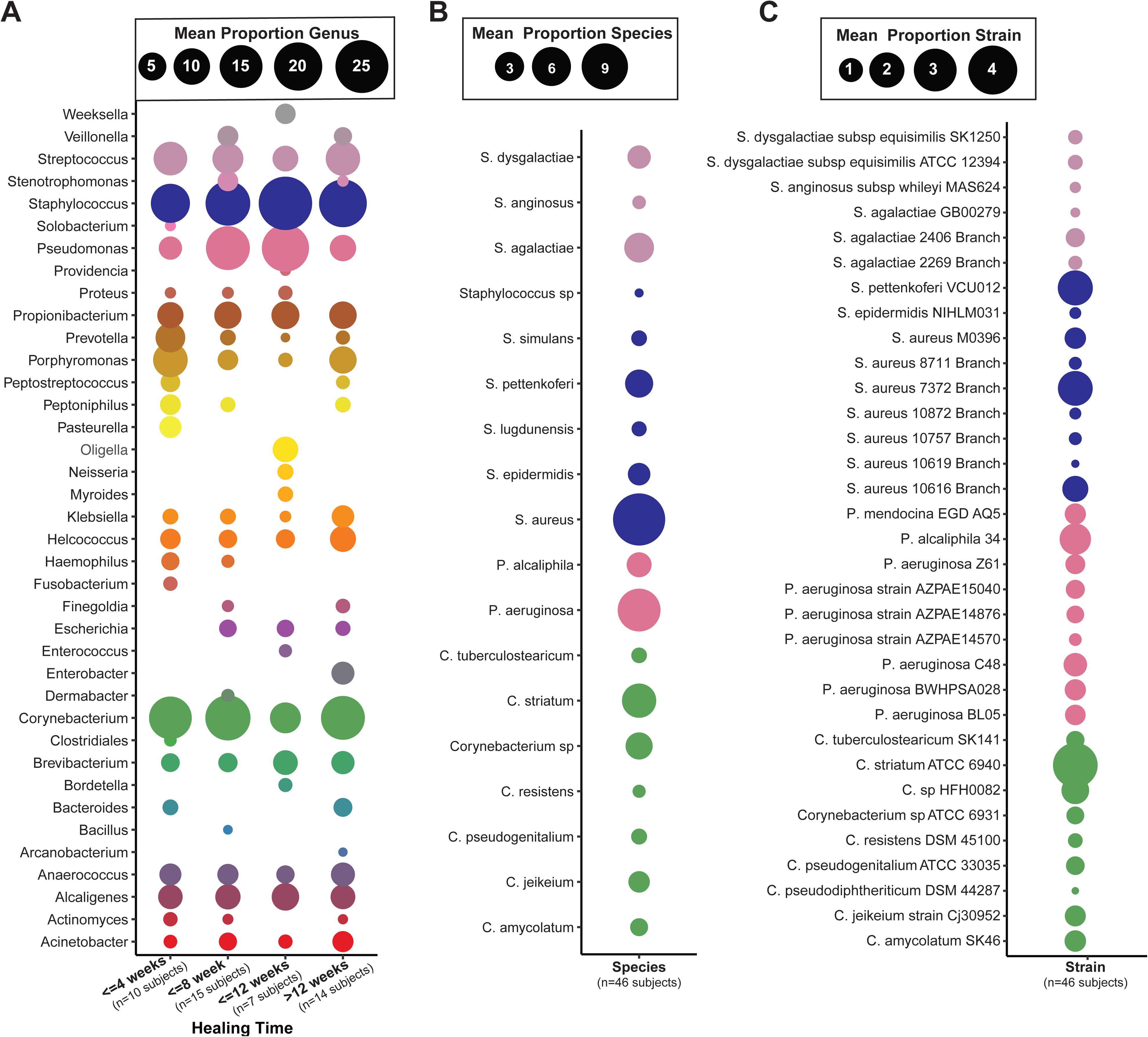
Strain-level resolution of DFU microbiota. A) Mean relative abundance of genera detected in >0.5% of samples from wounds with different healing rates. B) Most abundant bacterial species detected in >0.5% mean relative abundance from all samples of top genera. C) Most abundant bacterial strains detected in >0.5% mean relative abundance in all samples of top genera. Circle color indicates the taxonomic assignment; Circle size represents mean relative abundance.

### Effects of therapeutic interventions on DFU microbiota

Although antibiotic use is not indicated for DFU in the absence of overt clinical infection (heat, edema, purulent discharge)(Lipsky et al., 2012), 30 percent of our cohort (and 37 percent in our subset of 46 patients) were still administered systemic antibiotics during the follow-up period for wound-related complications or for other conditions that warranted antibiotic treatment (e.g. upper respiratory infection). We first tested the hypothesis that administration of systemic antibiotics selects for antimicrobial resistance in the DFU microbiota. To determine the prevalence of antibiotic resistance genes in DFU microbiota, we first examined the frequency of resistance to individual antibiotic chemical classes detected in each sample. At baseline, resistance to at least one class of antibiotic was detected in each subject, and genes conferring resistance to up to 10 different classes of antibiotics were detected in some wound specimens (Fig. 3A). Of antibiotic resistance classes detected, the most widespread were genes conferring resistance to beta-lactams, aminoglycosides, and macrolide antibiotics (Fig. 3B). In the thirty patients receiving antibiotic therapy, the resistance genotype did not correlate to the type of antibiotic administered (Supplementary Fig. 3). For example, cephalosporin administration does not drive an increase in the prevalence of beta-lactamase genes, although we note beta-lactam resistance is highly prevalent across the cohort, detected in 67% of all samples.

**Figure 3:**
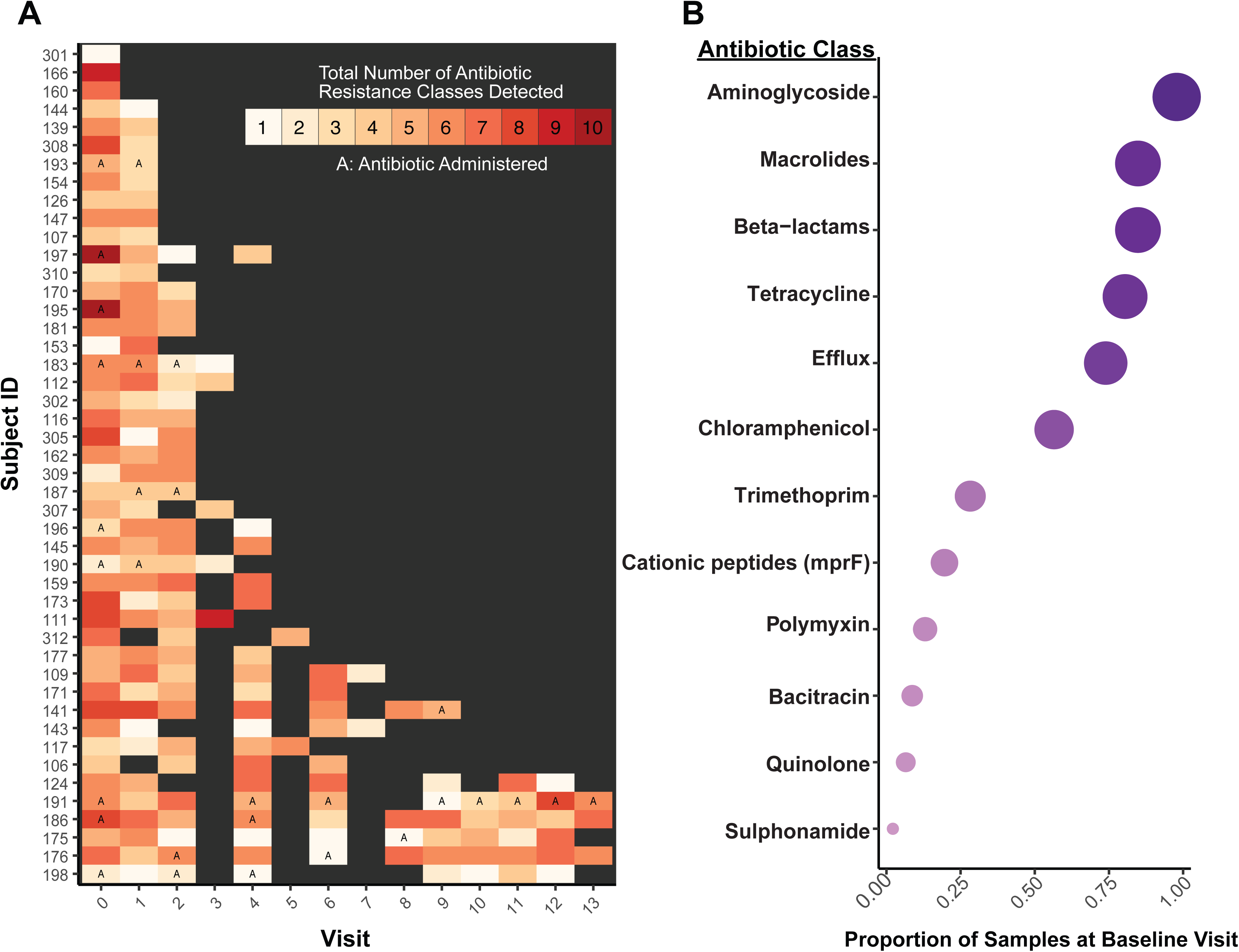
The DFU microbiome is multi-drug resistant. A) Timeline of each subject where the x-axis denotes the visit and the y-axis denotes individual subject IDs. The color of each visit corresponds to the total number of antibiotic resistance classes detected, with increasing darkness in red indicating increasing number of resistance classes detected. Grey boxes indicate a visit where the sample was either not sequenced, the wound was healed, or no resistance genes were detected. Visit 3, 5, and 7 were not sequenced unless it was the last visit a sample was collected before healing was recorded. Types of antibiotics with multiple classes of resistance (*e.g*., beta-lactamase class A, B, C etc.) were collapsed into a single class (*e.g*., beta-lactamases). The letter ‘A’ indicates a visit where antibiotics were administered. B) The proportion of samples with resistance genes detected (x-axis) for different classes of antibiotics (y-axis) at the baseline visit. Circle size corresponds to mean proportion.

The majority of DFUs that were not healed by week 12 remained unhealed or resulted in an amputation. Therefore, we used this time point to divide the cohort into ‘healing’ and ‘non-healing’ wound types, and to test the hypothesis that intervention differentially influences the microbiome of healing and non-healing wounds. Antibiotic administration did not have a significant effect on DFU microbiomes as measured by overall changes in alpha diversity before, during, or after the intervention in both groups (Fig. 4A). Since all subjects underwent sharp surgical debridement between specimen collection at the baseline study visit and the next study visit (t = 2 weeks), we also compared how this intervention, designed to remove necrotic tissue, influences wound microbiota. We determined that a significant reduction in Shannon diversity occurred in the visit immediately after debridement, but only in wounds that healed by 12 weeks (Fig. 4B). This suggests that microbiomes of non-healing DFUs did not respond to the initial debridement. When we examined the composition of the community at these time points, we observe that the relative abundances of aerobic bacteria such as *Staphylococcus, Streptococcus*, and *Pseudomonas* spp. do not change, however, mixed anaerobic bacteria such as *Anaeroccocus, Porphyromonas, Prevotella*, and *Veillonella* spp. are reduced after debridement in healed wounds, but not unhealed wounds (Fig. 4C,D).

**Figure 4:**
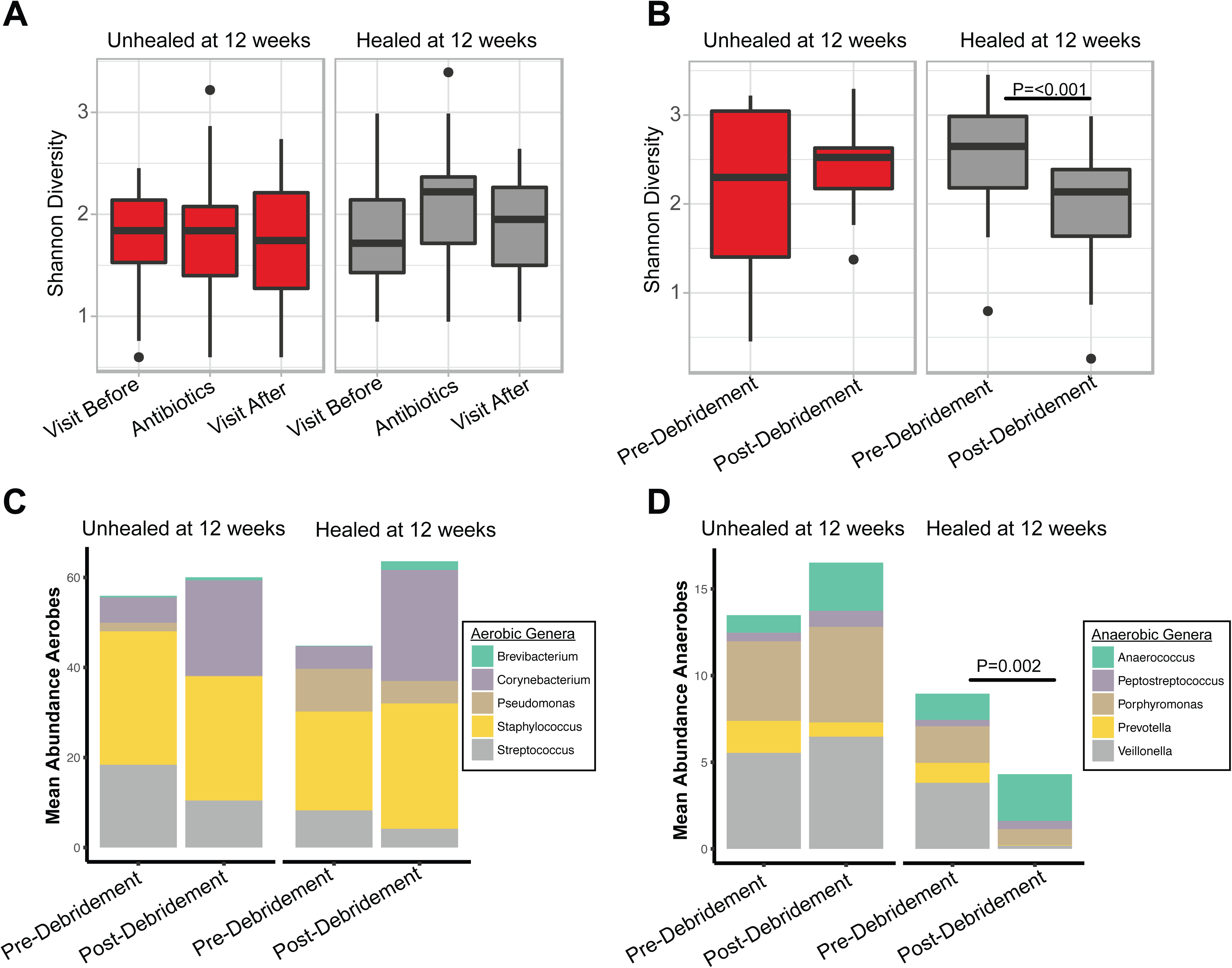
The DFU microbiome’s response to intervention predicts healing time. A) Shannon diversity remains unchanged in samples before, during, or after antibiotic administration in healing (n=9 subjects) and unhealed wounds (n=9 subjects). B) Debridement significantly reduces Shannon diversity in wounds that heal (n=32 subjects) within 12 weeks postdebridement. P<0.001, with non-parametric Wilcoxon rank-sum test. In wounds unhealed at 12 weeks (n=14 subjects) post-debridement a change in Shannon diversity is not observed. C) The mean proportion of common aerobic genera do not shift after debridement. D) The mean proportion of anaerobic genera are significantly reduced after debridement in wounds that heal within 12 weeks. P=0.002, with non-parametric Wilcoxon rank-sum test. In wounds unhealed at 12 weeks post-debridement the mean proportion of anaerobic genera does not change.

### Biofilm lifestyle is enriched in deep and poorly oxygenated wounds

To analyze gene functions of the DFU microbiome, we used the SEED database(Overbeek et al., 2005) to annotate microbial pathways. We then determined the relative abundance of SEED functions within each sample. We first took a high-level view of the top SEED subsystem level 1 functions within the dataset. The most abundant features included expected metabolic activities such as carbohydrate utilization, amino acid and protein metabolism. Also within the top functional roles are genes related to virulence, disease and defense, phages and transposable elements (Fig. 5A). After sub-setting the most abundant features in the dataset, we assessed correlations with clinical co-variates. Hierarchal clustering analysis of the resulting Spearman rank coefficients for SEED subsystem level 3 annotations (detected at >0.1% abundance) revealed that wound depth, surface area, and tissue oxygenation are correlated with differing microbial functional profiles (Fig. 5B). Deep and poorly oxygenated wounds are strongly associated with virulent metabolism, including capsular and extracellular polysaccharide production, saccharide biosynthesis, and non-glycolytic energy production (Fig. 5B). Associations for all SEED annotations can be found in Supplementary Fig. 4. Taken together, our data are suggestive of depressed metabolic activity and heterogeneity, hallmarks of an established biofilm, and further supports our hypothesis that in stable, non-healing wounds the microbiome exists in a biofilm state.

**Figure 5:**
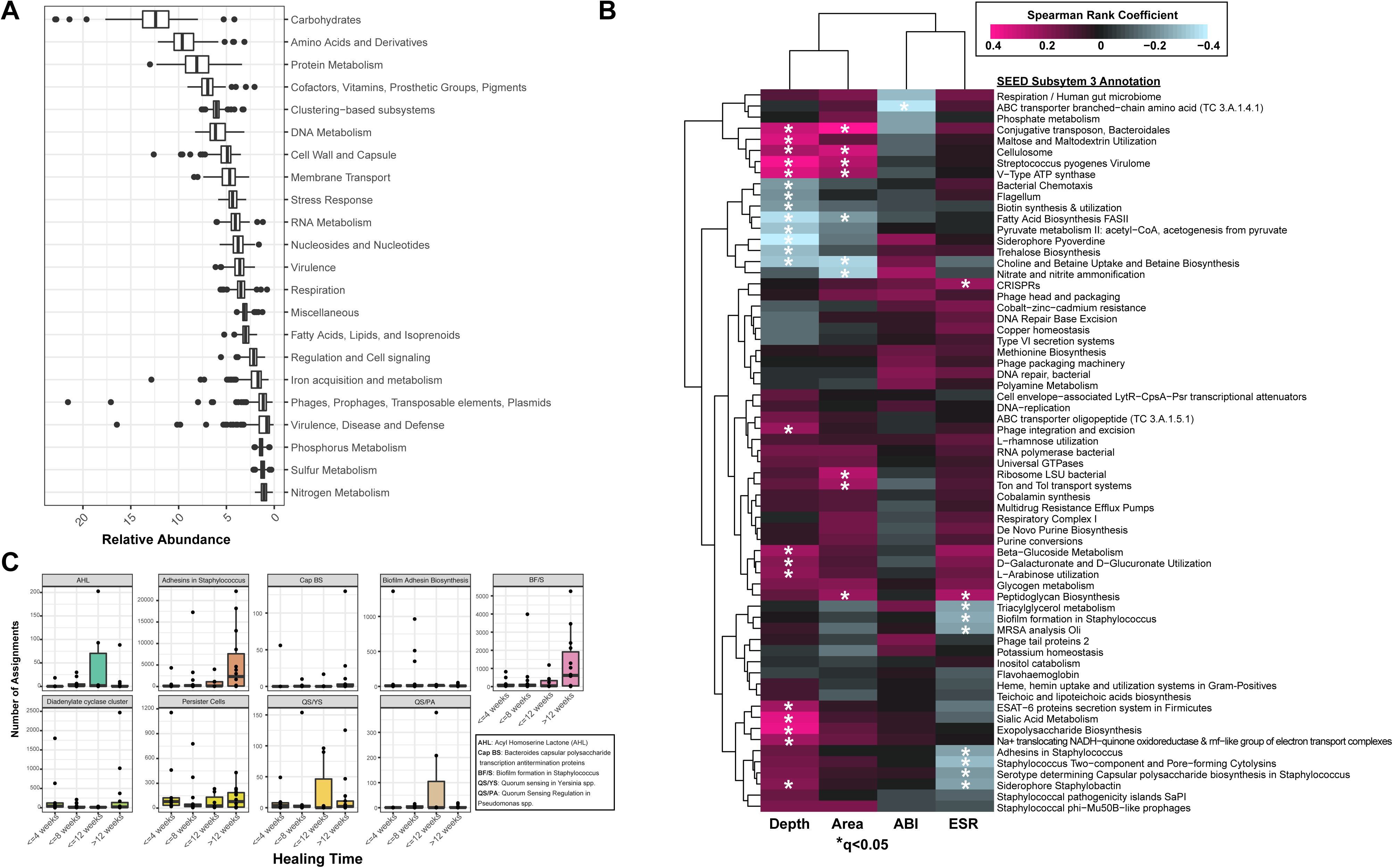
Metagenome annotation reveals functional subsystems associated with clinical factors and outcomes. A) Mean relative abundance of the top SEED subsystem level 1 annotations detected in DFU metagenomes. B) Correlation heatmap and hierarchical clustering of SEED subsystem level 3 annotations with clinical co-variates. Color corresponds to Spearman rank coefficient (pink and blue indicating positive and negative correlation, respectively). Wound depth and area cluster separately from ankle brachial index (ABI) and eosinophil sedimentation rate (ESR), markers of inflammation. Asterisk indicates significant associations (q<0.05). C) Number of read assignments to SEED subsystem level 3 annotations, normalized by total read depth per sample, with biofilm-specific terms (y-axis). Samples are stratified by healing time (x-axis). Each plot represents a single term.

*In situ* detection of biofilm is not clinically feasible without tissue biopsy and specialized microscopy techniques. Therefore, we elected to apply an indirect measurement of biofilm by extracting SEED annotations with terms related to biofilm formation such as ‘adhesins’, ‘biofilm’, and ‘persister cells’. We assessed the number of reads mapping to these functions by healing category, normalized by total read depth per sample. Slow or non-healing wounds were enriched in biofilm-related functional categories, compared to wounds that achieve closure by 12 weeks (Fig. 5C). We note that biofilm formation has been characterized in very few of the diverse microorganisms established to reside in wound tissue; thus those pathways are inherently excluded from this specific analysis and make our results even more compelling.

### *Staphylococcus aureus* strain diversity is associated with clinical outcomes

Mixed communities of Gram-positive cocci (GPC) are common to DFU. The GPC *Staphylococcus aureus* is an important skin pathogen due to virulence resulting in soft-tissue infection and emergence of both nosocomial and community-acquired multi-drug resistant strains. Culture-based diagnostics are able to successfully identify *S. aureus* and provide susceptibility testing, while amplicon-based 16S rRNA gene sequencing can identify *Staphylococcal* spp. at a lower limit of detection than culture, but species identification is limited. However, both of these methods are unable to differentiate individual strains of *S. aureus* without additional testing and genome sequencing. Furthermore, culture and amplicon-based studies have been unsuccessful in linking the presence of *S. aureus* to outcomes. Using a generalized linear model, we observed a significant association between *S. aureus* community abundance and healing time (Fig. 6A). However, 94% of our samples were positive for *S. aureus* in >0.1% abundance, demonstrating its ubiquity in a wide range of tissue environments, including those undergoing re-epithelialization and eventual wound closure, stable stalled wounds, and those developing osteomyelitis.

**Figure 6:**
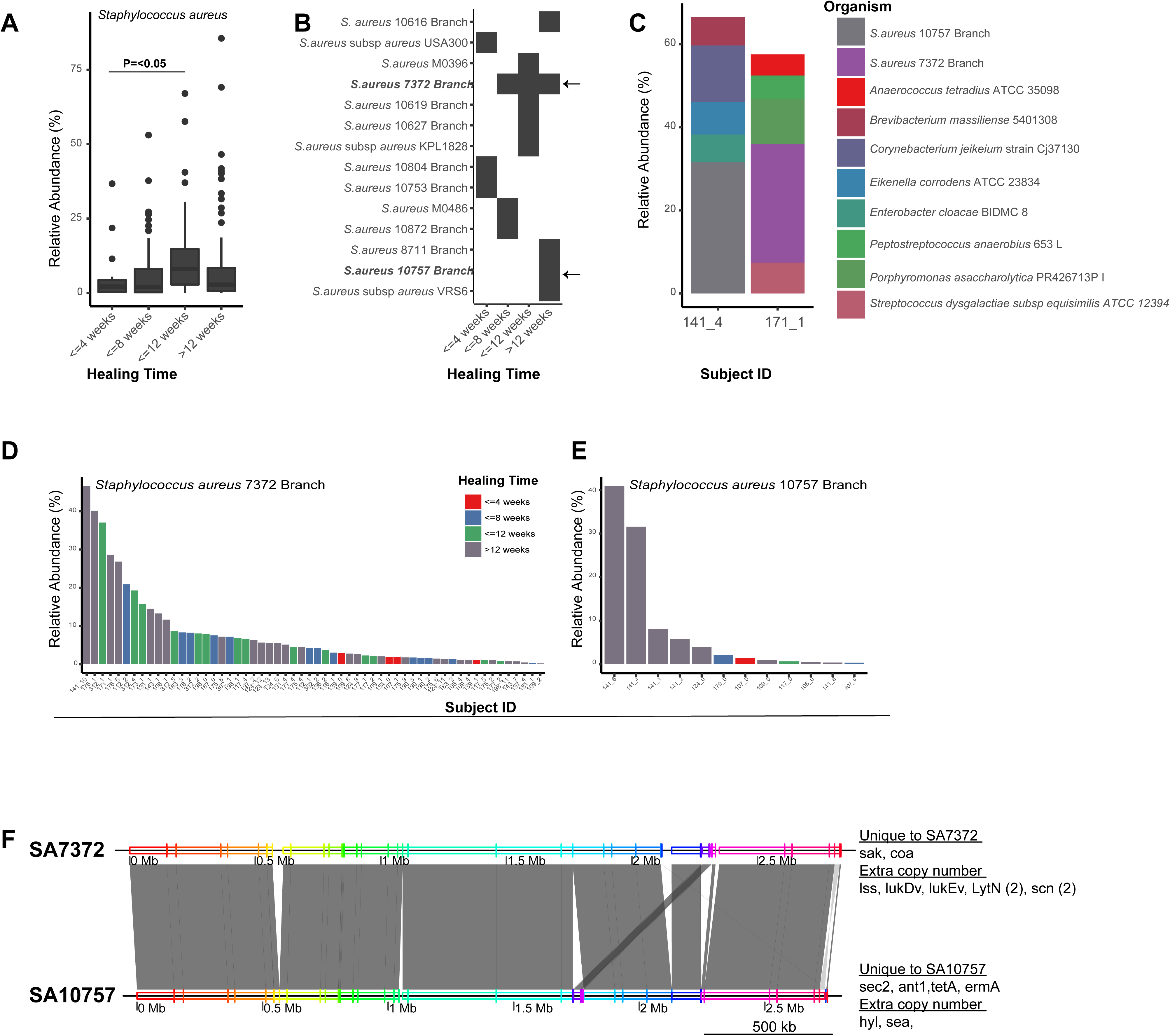
*Staphylococcus aureus* strain heterogeneity is associated with clinical outcomes. A) Mean relative abundance of *S. aureus* increases with healing time (P<0.05). B) Distribution of *S. aureus* strains and healing time. Each row corresponds to a different strain of *S. aureus* and the black box indicates detection in samples corresponding to each healing time (x-axis). Arrows indicate strains found in many samples (SA7372) and strains found only in non-healing wounds (SA10757). C) Microbiome community composition and taxa identified in >5 % relative abundance in patient specimens used to obtain representative isolates of SA7372 and SA10757. D) Mean relative abundance and distribution of SA7372 and E) SA10757 per sample across the cohort. Color indicates healing time. F) Whole genome alignment generated with MAUVE of SA7372 and SA10757. Grey shading indicates homologous blocks. Genes unique to each strain or found with extra copy numbers are denoted. SA7372 is enriched for virulence factors related to immune evasion (*e.g*., staphylokinase (*sac*), leucotoxins (*lukDv, lukEv*)). SA10757 is enriched for virulence factors related to antibiotic resistance and intoxication (*e.g*., hemolysin A (*hyl*), staphylococcal enterotoxin A (*sea*)).

We hypothesize that strain-level variation of *S. aureus* leads to variation in patterns of virulence and subsequent interference with tissue repair pathways. To test this hypothesis, we applied a phylogenetic approach to delineate species composition and identified unique strains of *S. aureus* present in wound tissue (Fig. 2C). In our cohort, some strains of *S. aureus* were broadly distributed across all healing categories. For example, *S. aureus* 7372 (SA7372) was detected in 28.7% (56/195) of DFU specimens across all healing times (Fig. 6B and D). We also identified several strains of *S. aureus* that are exclusively associated with unhealed wounds, such as *S. aureus* 10757 (SA10757), detected in 6.2% (12/195) of all specimens corresponding to 18.2% of non-healing wound specimens (Fig. 6B and E). These two representative strains of ‘generalist’ or ‘specialist’ *S. aureus*, respectively, are members of mixed GPC and anaerobic communities (Fig. 6C).

Because our findings associated *S. aureus* at the strain level with healing outcomes, we hypothesized that *S. aureus* strains vary in the degree to which they disrupt tissue repair. Furthermore, we hypothesize that clinical isolates of *S. aureus*, conventional wound pathogens, would elicit more severe host responses than isolates recovered from wounds that are generally regarded as non-pathogenic skin or environmental contaminants. To this end, we recovered SA7372, SA10757, and representative isolates of *Corynebacterium striatum* and *Alcaligenes faecalis*, to represent strains that are typically considered opportunistic pathogens but are not regularly identified in the clinical laboratory (C. *striatum*) and non-pathogenic environmental contaminants (A. *faecalis*)*. C. striatum* and *A. faecalis* were also the third and fourth most abundant species detected in our cohort, respectively (Fig. 1D, Supplementary Fig. 5A). We chose two specimens with greater than 10% abundance of SA7372 or SA10757 and used these samples to obtain pure cultures of *S. aureus* for further analysis (Fig. 6E).

To identify the genomic basis for eliciting differential host responses and clinical outcomes, we performed whole genome sequencing and comparative analysis on the two *S. aureus* isolates. The shared genome between the two strains consists of 2468 predicted genes comprising 90% of predicted open reading frames. Both strains contained the *agrABCD* operon encoding genes for the AGR quorum sensing system to produce autoinducing peptide (AIP) that functions to regulate biofilm development and virulence factors including toxins and degradative exoenzymes(Le and Otto, 2015; Novick and Geisinger, 2008).The genome of SA7372 contained 183 unique genes while the genome of SA10757 contained 64 unique genes. The majority of unique genes within each genome were of unknown function and predicted to encode hypothetical proteins. Staphylokinase (*sak*) is present in the SA7372 genome but not SA10757. In addition, SA7372 has an additional copy number of the genes encoding a neutrophil targeting leukotoxin (*lukDV*, lukEv)(Yoong and Torres, 2013), two extra cell wall hydrolases (*lytN*) that aide in protection from opsonophagocytic clearance(Becker et al., 2014), and two extra copies of the *scn*, encoding the Staphylococcal compliment inhibitor protein SCIN. Genes conferring resistance to aminoglycoside (*ant1*), tetracycline (tetA), and macrolide (*ermA*) antibiotics were unique to the SA10757 genome, in addition to the staphylococcal enterotoxin C-2 (*sec2*) and enterotoxin A (sea) (Fig. 6F).

To identify if these unique virulence related genes are present on mobile genetic elements, we used PHASTER to identify prophage elements within each genome. Staphylococcal enterotoxins *sec2* and *sea* were both present on an incomplete and thus likely defective phage element closely related to phage PT1028, suggesting stable integration in the SA10757 genome. An intact phage genome closely related to staphylococcus phage 96 containing the leukotoxin, hydrolase, and compliment inhibitor genes was detected within the SA7372 genome. Together, these findings provide evidence that *S. aureus* generalist and specialist strains differ at the genomic level in a phage-dependent manner to govern virulence-associated loci. This type of genomic diversification in turn may serve to influence host response and clinical outcome.

### The differential influence of primary wound isolates on host response and wound healing

To establish the functional implications of our metagenomic and genomic findings, we compared SA7372, SA10757, *A. faecalis*, and *C. striatum* clinical isolates using *in vitro* and *in vivo* approaches to assay host responses including wound healing. Because our data support the dogma that impaired tissue repair and prolonged ulceration are associated with the formation of a biofilm, we first tested the ability of each clinical isolate to form a biofilm *in vitro*. To better mimic the wound environment, we grew the biofilms on sterile cotton gauze, using keratinocyte culture media as the primary nutrient source. Each clinical isolate readily attached to and developed biofilms on individual cotton fibers of the gauze (Supplementary Fig. 5B).

To determine if wound isolates differentially influenced cytokine production by keratinocytes, we grew wounded primary keratinocytes in 1% cell-free spent media (CFSM) from mid-log phase planktonic bacterial cultures, or mature 72-hour biofilm cultures. After 8 hours of exposure, we quantitated the secretion of twenty inflammatory cytokines and applied hierarchal clustering to the resulting cytokine profiles (Supplementary Fig.6). We did not observe detectable levels of EGF, IFNR, IFN-α2, IL-1β, IL-2, IL-3, IL-4, IL-5, IL-7, or IL-9. Keratinocytes treated with *A. faecalis* biofilm and planktonic CFSM (Supplementary Fig. 6) exhibited a strong IL-8 response (1873 ± 202 AF biofilm vs. 36±12 pg/mL control; P=<0.0001). Further, *A. faecalis* biofilm and to a lesser extent planktonic CFSM resulted in a significant and specific increase in the production of G-CSF, GM-CSF, IL-6, TGF-α, TNF-α, and IP-10 compared to the control and other treatment groups (Fig. 7A). *A. faecalis* biofilm and *C. striatum* planktonic CFSM enhanced the production of platelet derived growth factor (PDGF-AB:BB,) while *C. striatum* planktonic CFSM was associated with an increase in IL-1α and IL-1RA. Exposure to *S. aureus* CFSM from biofilm or planktonic cultures did not significantly shift cytokine production levels compared to the untreated control group, with the exception of SA7372 biofilm CFSM which resulted in decreased production of TGF-α (Fig. 7A; Supplementary Figure 6). This trend was also observed for *C. striatum* biofilm and planktonic CFSM, whereas *A. faecalis* CFSM resulted in an increased production of TGF-α.

**Figure 7:**
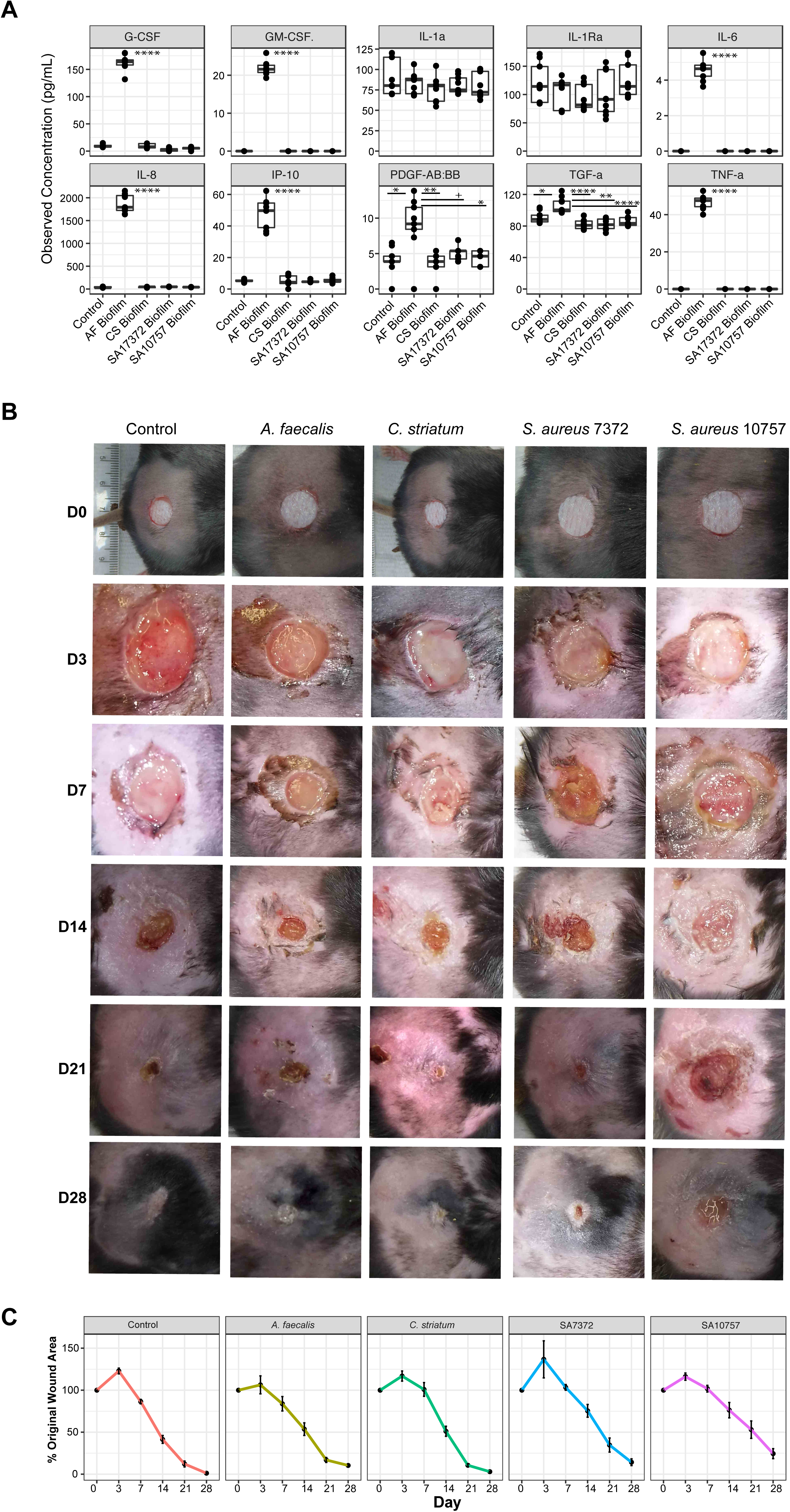
Primary wound isolates result in differential host responses and wound healing. A) Observed concentration of secreted cytokines (pg/mL) from primary keratinocytes exposed for 8 hours to conditioned media from mature biofilms of *A. faecalis* (AF), *C. striatum* (CS), *S. aureus* 7372 (SA7372), or *S. aureus* 10757 (SA10757). Each condition was repeated with three biological replicates and three technical replicates of each (n=9). Analysis of variance with post hoc multiple comparison testing was performed between each group (****<0.00001, ***<0.0001, **<0.001, *<0.01, +<0.05). B) Biofilms of each strain listed above were allowed to mature over a period of 72 hours on sterile gauze before being placed into full thickness dorsal mouse wounds (n=4 mice per group). Photographs of the wounds were taken at day 0, 3, 7, 14, 21, and 28. Wound measurements were recorded by two independent observers and are plotted in C) over time. Error bars represent standard error of the mean.

To determine if different strains of primary clinical isolates impact healing rates *in vivo*, we used a type II diabetic mouse model of impaired wound healing (db/db; *Lepr^−/−^*). These experiments were performed with mature biofilms since the most pronounced effects on keratinocyte cytokine expression occurred when treated with biofilm, and because we hypothesize that within wound tissue bacterial isolates grow as biofilms. Full thickness excisional wounds were created on the mouse dorsa using 6 mm punch biopsy and mature biofilms were transferred into the wounds. A non-infected negative control consisted of gauze soaked in PBS. Each wound was photographed and measured on days 0, 3, 7, 14, 21 and 28 (Fig. 7B). Average wound surface area (percent of the original wound size), increased in all groups by day 3, except *A. faecalis* (ctrl = 123 %, C. striatum = 117 %, SA7372 = 137 %, SA10757 = 116 %). Notably the wound margins in the *A. faecalis* group remained defined while they became irregular, diffuse, and macerated in the other infection groups, including the non-infected control. By day 7, wounds infected with *A. faecalis* biofilm resumed the same healing-trajectory as the control group (Fig. 7C). The *C. striatum* group exhibited an early delayed healing phenotype, with a mean wound area of 100.9 % of the original area on day 7, compared to 86.4% mean wound area in the control group. However, by day 14 the *C. striatum* infected wounds resumed the control healing trajectory. Persistent delayed healing occurred with both clinical *S. aureus* strains. By day 21 the control group exhibited near complete closure (mean wound area percentage of original=12.1%), while the mean original percent wound area of wounds infected with SA7372 and SA10757 was 34.6% and 53.1% respectively (P=0.01 and P=0.005). On day 28, the wounds inoculated with the strain of *S. aureus* detected in unhealed wounds by metagenomics, SA10757, exhibited the slowest healing rate and open wounds remained with a mean percent wound area of 24% of the original size. Together, these findings demonstrate the differential influence of *S. aureus* strain-level variation and other primary wound isolates on healing *in vivo* and provide functional evidence for the microbial basis of delayed healing in chronic wounds.

## DISCUSSION

Diabetic foot wounds are complicated by several factors that contribute to impaired tissue regeneration including hyperglycemia, peripheral neuropathy, vascular disease and a complex microbiome. Microbial communities that assemble in wound tissue are difficult to detect and are not necessarily associated with cardinal signs of infection(Lipsky et al., 2012), further complicating prognostics for wound healing outcomes. Chronic wounds have a major societal impact; thus our in-depth investigation of the DFU microbiome, coupled with *in vitro* and *in vivo* functional modeling, enhances our understanding of microbial influences on tissue repair pathways, suggests new diagnostic/prognostic and therapeutic targets, and has the potential to overcome challenges for improving patient outcomes.

Here we apply shotgun metagenomic sequencing of time-series specimens from patients with DFU to achieve strain-level classification of microbial communities. This allowed us to pursue guided isolation and whole genome sequencing of specific microbial strains from matched patient derived samples. Despite the growing body of literature dedicated to the study of wound microbiomes, all studies to date have exclusively employed amplicon-based sequencing of phylogenetic marker genes such as the 16S rRNA gene, failing to distinguish individual species. For example, within the *Staphylococcus* genus are skin commensals and the notorious pathogen *S. aureus*. Clinically, *S. aureus* would be regarded differently than the commensal *S. epidermidis;* thus their classification is critical for determining efficacious treatment strategies. Additionally, shotgun metagenomics allows for strain-level tracking and functional annotation, which both revealed novel aspects of the DFU microbiome and its association with clinical outcomes in this study.

The inherent properties of a diabetic wound environment support the establishment of a diverse microbiome, although antibiotic use is not recommended for DFU in the absence of overt clinical infection, such as cellulitis and osteomyelitis(Lipsky et al., 2012). However, some cohorts have reported antibiotic use in 60% of patients(Howell-Jones et al., 2006; Siddiqui and Bernstein, 2010; Tammelin et al., 1998), potentially selecting for antibiotic resistance, and it was a DFU from which the first strain of vancomycin-resistant *S. aureus* was isolated(Tenover et al., 2004). We characterized the patterns of antibiotic resistance both across our cohort and over time to determine that antibiotic resistance is widespread and DFU microbiomes are multi-drug resistant, in some cases harboring genes conferring resistance to ten different classes of antibiotics. Greater than 50% of wound specimens contained resistance genes to the aminogylocoside (e.g., clindamycin), macrolide (e.g., erythromycin), beta-lactam (e.g., amoxicillin), and tetracycline (e.g., minocycline) classes of antibiotics. We further examined the effects of antibiotic administration at the community level by measuring changes in alpha diversity. We determined that antibiotics do not change the overall diversity in healed or non-healed wounds, suggesting little perturbation to the microbiome within the wound. Our findings offer further support toward guidelines to reserve the use of antibiotics for clinically overt infections and suggest their use does not positively impact the overarching goal of wound healing.

It is recommended that all diabetic wounds are surgically sharp debrided, to remove debris, callus, necrotic (senescent, devitalized), and infected tissue(Lipsky et al., 2012). This procedure is thought to ‘re-activate’ stalled healing pathways by inducing an acute wound(Ashrafi et al., 2016). Although correlated with improved healing rates, the association is not significant and less than half of debrided DFUs progress towards healing(Cardinal et al., 2009). To identify microbiome signatures that could differentiate wounds that respond positively to debridement, we used Shannon’s diversity to assess global shifts in the community structure pre- and post-debridement. The microbiome of healing wounds, defined by complete re-epithelialization achieved within 12 weeks of debridement, exhibited marked decreases in diversity, driven by a reduction in the abundance of anaerobic bacteria in the overall community. Compromised blood flow leads to local tissue ischemia that can promote growth of anaerobic microorganisms. Several targeted studies have concluded that anaerobes are underrepresented in culture-based estimation of DFU isolates(Citron et al., 2007; Louie et al., 1976). We conclude several species of anaerobic bacteria are abundant across DFUs in association with mixed aerobes, and our data further suggest that successful debridement is concomitant to disrupting anaerobic networks. Thus we propose that the microbiome can serve as a prognostic marker of healing trajectory at the time of debridement in order to target DFUs without decreases in microbiome diversity after debridement for more extensive therapy.

Given that *Corynebacterium* was the second most abundant genera classified in our cohort by both 16S rRNA gene sequencing and shotgun metagenomics, consistent with previous chronic wound microbiome studies(Dowd et al., 2008; Gardner et al., 2013; Loesche et al., 2017; Rhoads et al., 2012; Wolcott et al., 2016), we hypothesized that *Corynebacterium* spp. have a more significant role in DFU than simple contamination from intact skin. We found that standard clinical microbiology protocols do not routinely classify *Corynebacterium* spp. without the use of specialized workflows, but instead group aerobic Gram-positive, catalase-positive rods as diphtheroid and consider them to be skin contaminants(Leal et al., 2016). *Corynebacterium striatum* is the most prevalent and abundant *Corynebacterium* spp. in DFU, detected in 28% of our specimens. Considered an emerging multi-drug resistant microbe(Hahn et al., 2016), *C. striatum* is an underrecognized cause of diabetic foot osteomyelitis(G.J. et al., 2010; Patel et al., 2016) suggesting it should not be classified as merely contaminating skin flora in the clinic. *Brevibacterium massiliense* is another diphtheroid detected in 56 of 195 specimens, often in >10% relative abundance of the community. Little is known about this species of Actinobacteria that was first isolated from wound discharge and subsequently classified in 2009(Roux and Raoult, 2009). We note that in this study, the isolate was first identified as *a Corynebacterium* species by biochemical testing until molecular characterization confirmed it was within the genus *Brevibacterium*. Thus, *B. malssiliense* is just one example of a bacterial species we have found to be more widespread in DFU than previously reported, whose role in the context of wound healing is undefined.

To investigate the biological consequences of *S. aureus* strain variation and colonization with microbes commonly dismissed as skin flora (*C. striatum*) or environmental contamination (*A. faecalis*), we measured differences in the inflammatory response in primary keratinocytes cultured in biofilm and planktonic media of primary wound isolates. We found the environmental Gram-negative rod *A. faecalis*, normally considered nonpathogenic, induces a striking keratinocyte response via production of pro-angiogenic/pro-inflammatory cytokine IL-8(Rennekampff et al., 2000),(Arwert et al., 2012) in addition to cytokines known to stimulate proliferation and enhance wound healing (GM-CSF, G-CSF, PDFG-AB). Diabetic wounds with *A. faecalis* biofilms healed at an accelerated rate during the early stages of wound healing. These findings suggest a beneficial role for some microbes in tissue repair. Future studies should address how *A. faecalis, C. striatum* and other “non-pathogenic” wound microbiota function in a polymicrobial setting to influence virulence of wound pathogens, host responses, and healing outcomes.

Strain heterogeneity of *S. aureus* is associated with disease severity in other dermatological conditions such as atopic dermatitis. Particularly, strains from more severe patients elicit stronger immune responses and skin inflammation(Byrd et al., 2017). Diabetic wounds are consistently colonized by *S. aureus* so we focused our analysis on this skin pathogen to classify strains associated with different wound healing outcomes. We identified *S. aureus* strains with a wide host range and strains exclusively associated with unhealed wounds. The genome of *S. aureus* associated with poor wound healing outcomes harbored multiple antibiotic resistance genes and genes encoding staphylococcal enterotoxins. These superantigens result in an exacerbated inflammatory response by non-specific stimulation of large populations of T-cells(Ortega et al., 2010), suggesting colonization by these strains results in persistent inflammation leading to impaired healing progression. Conversely, genome analysis of *S. aureus* isolates with non-specific associations suggest such strains are experts at immune evasion and warrant additional investigation to link genome diversity with phenotypic differences in pathogenesis.

Chronic wounds are a major strain to health care systems and cause significant morbidity and mortality. As the rate of diabetes and obesity increases worldwide, the economic and social burden of chronic wounds such as DFU is projected to snowball(Sen et al., 2009). Therefore, new approaches for their management and treatment are desperately needed. Here, we applied shotgun metagenomic sequencing to a common type of chronic wound to identify microbial taxonomic and genetic markers associated with clinical outcomes. By coupling metagenomic and genomic analyses with *in vitro* and *in vivo* models of host response and wound repair, we demonstrate the functional implications of a) strain-level variation of the wound pathogen *S. aureus* and b) commonly disregarded isolates such as skin commensals. Our findings suggest new targets for diagnostic and prognostic markers to guide treatment and a potential for targeted microbial-based therapeutics to improve healing and clinical outcomes.

## ACKNOWLEDGMENTS

**Funding:** The National Institutes of Health, National Institute of Nursing Research (R01-NR-009448 to S.E.G, R01-NR-015639 to E.A.G, and P20 NR018081 to SEG), National Institute of Arthritis, Musculoskeletal, and Skin Diseases (R01-AR-006663 and R00-AR-060873 to E.A.G.), Burroughs Wellcome Fund PATH Award (E.A.G), Linda Pechenik Montague Investigator Award (E.A.G.). This research was also supported by the Penn Skin Biology and Disease Resource-based Center (P30-AR-069589). M.L. was supported by the Dermatology Research Training Grant, T32-AR-007465. We would also like to acknowledge Gordon Ruthel and the Penn Vet Imaging Center for support with confocal microscopy.

---

**Supplementary Figure 1:** Alpha diversity metrics of the DFU microbiome for 16S rRNA gene sequencing versus shotgun metagenome sequencing. A) Shannon Index and B) Observed Number of Species

**Supplementary Figure 2:** Most Abundant Taxa Identified in Diabetic Foot Ulcers. A) Isolates cultured in the laboratory from DFU. Proportion indicates total proportion of all isolates (n=1245 from a total of 384 samples). B) Taxa identified by 16S rRNA gene sequencing of the V1-V3 hypervariable region from the same samples and DNA extracts as this study. Mean proportion calculated as mean proportion in all samples >1%.

**Supplementary Figure 3:** Antibiotic resistance patterns in patients administered systemic antibiotics. Each box represents a single patient labelled by the class or classes of antibiotic administered. The visit before, visit of, and visit after antibiotics are given is along the x-axis and each class of antibiotic resistance detected is plotted along the y-axis and represented as a (*) to indicate positive detection.

**Supplementary Figure 4:** Functional SEED subsystems level 3 associated with all clinical factors and outcomes. Shown is a correlation heatmap and hierarchical clustering of SEED subsystem level 3 annotations with all measured clinical co-variates. Color corresponds to Spearman rank coefficient (red and blue indicating positive and negative correlation, respectively).

**Supplementary Figure 5:** Biofilm formation by wound isolates. A) Mean proportion of each species targeted for *in vitro* and *in vitro* characterization by healing time. B) Representative image of biofilm formation by each isolated species and stained with the LIVE BacLight Bacterial Gram Stain Kit.

**Supplementary Figure 6:** Detection of cytokines from primary keratinocytes exposed for 8 hours to conditioned media from mature biofilms or mid-log phase planktonic cultures of *A. faecalis* (AF), *C. striatum* (CS), *S. aureus* 7372 (SA7372), or *S. aureus* 10757 (SA10757). Each condition was repeated with three biological replicates and two or three technical replicates of each (n=6-9). A) Heatmap and hierarchal clustering of all conditions displaying the mean concentration of cytokines measured (pg/mL). B) Observed concentration of secreted cytokines (pg/mL) after exposure to planktonic cultures.

## MATERIALS AND METHODS

### Study Design

The study design, subject enrollment, and specimen collection are described in previous publications(Kalan et al., 2016; Loesche et al., 2017). Briefly, 100 subjects were enrolled in a prospective-cohort to sample the DFU microbiota and measure outcomes. Samples for microbiota analyses were collected at initial presentation (V0) and every two weeks until the DFU: i) healed; ii) was amputated; or iii) 26 week of follow up elapsed (V1-12). The Institutional Review Boards at the University of Iowa (IRB#200706724) and the University of Pennsylvania approved the study procedures (IRB#815195). Informed consent was obtained from all participants in writing.

Wound management was standardized to aggressive sharp debridement of necrotic tissue in the wound bed at baseline and wound edge callus removal every two weeks followed by non-antimicrobial dressing application (i.e., Lyofoam^®^, Molnlycke Health Care). Ulcers were offloaded with total contact casts (87 subjects) or DH boots (13 subjects). Topical antimicrobial or system antibiotic administration was not included unless an infection-related complication was present at baseline or occurred within the study period. Data was collected at baseline and longitudinally every two weeks until the wound healed or 26 weeks elapsed for a total of 384 wound specimens. We subset this dataset to maximize read depth and output for shotgun metagenomic sequencing and this is described in the results section.

### Study Variables

*Clinical factors:* Demographic variables were collected at the baseline visit including age, sex, diabetes type and duration and duration of the study ulcer using subject self-report and medical records. At each study visit glycemic control was measured by levels of haemoglobin A1c and blood glucose. Inflammatory (Erythrocyte sedimentation rate (ESR), C-reactive protein) and immune (white blood cell counts) markers were determined with standard laboratory tests.

Each subject was also assessed for ischemia using the ankle-brachial and toe-brachial index and for neuropathy using the 5.07 Semmes-Weinstein monofilament test. Transcutaneous oxygen pressure was measured at baseline and at each follow-up visit, using a transcutaneous oxygen monitor (Novametrix 840^®^, Novametrix Medical Systems Inc.). Ulcer location was categorized as forefoot, midfoot, or hindfoot. The level of necrotic tissue was defined as black, grey or yellow tissue in the wound bed measured using a likert scale as the percentage of the total wound area binned according to 0-25%, 25-50%, 50-75% or 75-100% necrotic tissue.

*Outcomes:* Healing and infection-related complications were assessed every two weeks. Ulcer closure was determined using the Wound Healing Society’s definition of “an acceptably healed wound,”(Margolis et al., 1996). “Infection-related complications” included wound deterioration defined as development of frank erythema and heat, and an increase in size > 50% over baseline, new osteomyelitis, and/or amputations due to infection. Two members of the research team independently assessed each DFU for erythema and heat. Two members of the research team independently assessed size using the VeVMD^®^ digital software system (Vista Medical, Winnipeg, Manitoba, Canada) and procedures previously described(Gardner et al., 2012). A cotton-tipped swab, placed in the deepest aspect of the DFU, was marked where the swab intersected with the plane of the peri-wound skin. The distance between the tip of the swab and the mark was measured as ulcer depth using a centimetre ruler.

Osteomyelitis was assessed using radiographs and MRI at baseline and during follow-up visits when subjects presented with new tracts to bone, wound deterioration, elevated temperature, elevated white count, elevated erythrocyte sedimentation rate, or elevated C-reactive protein. If these indicators were absent at follow-up, radiographs were not retaken. Subjects experiencing new amputations had their medical records reviewed by the research team to ensure amputations were due to DFU infection, and not some other reason.

*Metagenomic Sequencing of DFU specimens:* After cleansing the ulcer with sterile saline, specimens were collected using the Levine technique by rotating a swab over a 1-cm^2^ area of viable non-necrotic wound tissue for 5 seconds with sufficient pressure to extract tissue fluid. DNA was isolated from swab specimens as previously described(Loesche et al., 2017). To minimize signal from contaminating eukaryote DNA (human), a microbial DNA enrichment step was performed prior to Illumina library preparation with the NEBNext Microbiome DNA Enrichment kit (New England Biolabs). Purified DNA was quantified using the Qubit dsDNA High-Sensitivity Assay Kit (Invitrogen). Illumina sequence libraries were prepared using the NexteraXT DNA Library Preparation Kit (Illumina) and followed by quality control (Bioanalyzer), quantification (Kapa assay), pooling, and sequencing at either the University of Maryland Institute for Genome Sciences or with CosmosID. 150 bp paired-end sequencing of pooled samples was performed over four runs of two full flow cells (16 lanes) on the HiSeq 4000 to generate 200-400M reads per sample.

The raw read data were first preprocessed in collaboration with CosmosID to filter and remove human DNA sequences by mapping reads to the human genome and a custom built database of human DNA sequences, followed by additional filtering for repeat regions using the Tandem Repeat Finder algorithm (https://tandem.bu.edu/trf/trf.html). Finally, filtered reads were mapped to custom curated bacterial, fungal, viral, and antibiotic resistance genomic databases. Taxonomic identification was assigned with an in-house K-mer based algorithm refined against a whole genome phylogenetic tree to identify unique species and strains developed at CosmoID and described in Hourigan et al(Hourigan et al., 2018).

Filtered reads were further processed with an in-house pipeline (Grice lab) that included additional read QC and linker cleaning steps using the cutadapt software(Martin, 2011). Microbial ecology metrics including Shannon diversity index and number of observed species (richness) were calculated in the R statistical environment using the vegan package(Oksanen et al., 2018; R core team, 2017). Functional annotation was assigned with the SUPERFOCUS software (Silva et al., 2015) which classifies each metagenomic sequence into a SEED subsystem(Overbeek et al., 2014) run in the sensitive mode, using the diamond aligner and db_98 with the following command: python SUPERFOCUS_0.27/superfocus.py -q. -m 1 -a diamond - fast 0 -o all_samples -t 24 -dir SUPER-FOCUS/

Antibiotic resistance gene classification was collapsed into functional categories (e.g., class A, B, and C beta-lactamases were grouped as “beta-lactamase”).

Raw data are under submission and pending accession number assignment in the NCBI Sequence Read Archive (SRA) under the BioProject Accession number PRJNA324668.

### Whole Genome Sequencing

DNA was isolated from pure culture following the same procedure as microbiome samples described above. Purified DNA was used as input to the NexteraXT library preparation kit following the manufacturers protocol (Illumina). Libraries were sequenced on the NextSeq500. Raw data was processed with cutadapt(Martin, 2011) to trim adapter sequences and a minimum quality score of 20. Reads were assembled with Unicycler(Wick et al., 2017) and annotated with Prokka(Albertsen et al., 2014; Seemann, 2014). Contigs were then ordered with CAR(Lu et al., 2014) prior to full genome alignment using MAUVE(Darling et al., 2004). The two *S. aureus* genomes were compared for shared and unique genes using Roary(Page et al., 2015) and default settings.

Phage sequences were detected by running genomic contigs through the PHASTER(Arndt et al., 2016) algorithm and classified as complete, questionable or incomplete.

### Bacterial Isolation and Manipulation

*Isolation:* Bacterial isolates were grown from wound swabs collected in trypticase soy broth (TSB) and described in Gardner et al. 2013. Identification was achieved with the Vitek Legacy (Biomerieux) or full length 16S rRNA gene Sanger sequencing.

*Planktonic and Biofilm growth:* Isolates were grown overnight at 37°C on TSA. Colonies were scraped into Keratinocyte media without antibiotics to an OD_600_ nm of 0.08-0.1. Keratinocyte media was made by mixing Keratinocyte SFM (kit 17005042, Life Technologies) with Medium 154 (M154500, Life Technologies) supplemented with human keratinocyte growth supplement (S0015, Life Technologies) at a 1:1 ratio

For planktonic cultures the inoculums were then added in a 1/10 dilution to a final volume of 5 mL of Keratinocyte media. Biofilms were inoculated in a 1/10 dilution to a final volume of 3 mL keratinocyte media in 6 well polystyrene plates containing a 1 inch × 1 inch piece of sterile cotton gauze. Cultures were incubated at 37°C for 72 hrs, shaking at 200 rpm to allow adhesion and growth, with the media changed every 24 hrs. After incubation, the media was removed and the biofilms attached to the gauze were washed with 5 × 1 mL sterile phosphate buffered saline to remove non-adherent cells. For imaging, the biofilms were stained with the LIVE BacLight Bacterial Gram Stain Kit (ThermoFisher Scientific, Waltham, MA) with SYTO9 (480nm/500nm) and hexidium iodide (480nm/625nm) as a 2 mL solution in water and according to the kit instructions. The stain was removed and deionized water was added to the biofilms prior to imaging on a Leica SP5-II inverted confocal microscope. Images were post-processed with Volocity software (PerkinElmer, Waltham, MA, USA). The maximum projection for each image was used to generate supplementary figure 5. Quantitative counts of either planktonic or biofilm cultures (after disruption by vortexing for 3 × 1 min each) was performed by serial dilution in phosphate buffered saline. The dilutions were plated onto non-selective media (TSA) and incubated for 16-18 hrs at 37°C. Colonies were counted and the total CFUs calculated.

*Keratinocyte Cytokine Analysis:* Primary keratinocytes were obtained from the University of Pennsylvania Skin Biology and Diseases Resource Center and cultured in keratinocyte media (see above) with 1% antibiotic/antimycotic (15240062, Life Technologies). Cells were seeded at 0.3 × 10^^^6 cells/well in a 6-well tissue culture plate (2.5 mL total volume per well). Media was changed the day after seeding, and then every other day until confluent. Confluence was achieved after ~4 days. Planktonic or biofilm cultures were prepared as described above except planktonic cultures were grown to mid-log phase. Based on quantitative counts of each culture, biofilm or planktonic cultures were normalized to a count of 2.8 × 10^7 cfu/mL, which was the lowest cell density of the four test groups *S. aureus* 10757, *S. aureus* 7372, *Alcaligenes faecalis*, and *Corneybacterium striatum*. Cultures were centrifuged to remove the majority of cells and filtered through a 0.2 uM filter for sterility. Each filtered planktonic culture was then diluted 1/50 and each filtered biofilm culture diluted 1/100 into keratinocyte media before being placed on the keratinocytes. Prior to exposure, a single ‘scratch’ was made across each cell monolayer with the a pipette tip to mimic a wound and then washed three times with sterile saline to remove cell debris. The keratinocytes were exposed to the filtered spent media from. biofilm or planktonic cultures for 8 hours. Three biological replicates of each condition was performed. Media was collected from each well into 1.5 mL microcentrifuge tubes, centrifuged at maximum speed for 10-15 min. The supernatant was filtered through a 0.2 uM filter and placed at −20 Celsius until analysis. Secreted cytokine concentrations were determined by Luminex Multiplex Assay Human Cytokine/Chemokine Panel I (MilliporeSigma) and was performed at the University of Pennsylvania Human Immunology Core (P30-CA016520). Two technical replicates of each biological replicate for mid-log phase treatments were performed (n=6 per treatment) and three technical replicates of each biological replicate were performed for the biofilm treated cells and control cells with no treatment (n=9 per treatment).

*Diabetic Mouse Wounding and Infection:* Twenty 8-wk-old female BKS.Cg-Dock^7m+/+^Lepr^db^/J mice were purchased from Jackson laboratories. All mice were housed and maintained in a BSL II and specific pathogen free conditions at the University of Pennsylvania under and approved IACUC protocol (804065). At 12-wks old a dorsal patch of skin was shaved and the mice were then housed individually for three days. On day 4, after anesthesia by inhaled isoflurane, an 8-mm full-thickness excisional wound was created with a punch biopsy tool (Miltek). Seventy-two hour biofilms grown on cotton gauze and precisely cut to 8-mm with a clean sterile biopsy tool were then placed into the wounds and covered with a transparent film (Tegaderm, 3M). Each piece of 8-mm size gauze contained between 1.23 − 9.26 × 10^^^8 cfu. Four mice per treatment group were included, the control group was treated with uninoculated sterile gauze treated in the same manner as the biofilm groups (described above). Each wound was photographed and measured at the time of wounding (t=0), day 3, day 7, day 14, and day 28 post-wounding. After day 3 the gauze was removed and the wound was recovered with the transparent film. Wound area was measured in Adobe photoshop by two independent persons and the average of each measurement recorded for each individual wound.

### Data Analyses

The R Statistical Package(R core team, 2017) was used to generate all figures and compute statistical analysis. Statistical methods are described within the text and figure legends. Correlations between SEED functional categories and clinical features were determined by calculating the Spearman coefficient.

